# ME-plot: A QC package for bisulfite sequencing reads

**DOI:** 10.1101/033696

**Authors:** Bong-Hyun Kim

## Abstract

**Summary:** Bisulfite sequencing is the gold standard method for analyzing methylomes. However, the effect of quality control in bisulfite sequencing has not been studied extensively. We developed a package ME-Plot to detect the errors in bisulfite sequencing mapping data and produce higher quality methylation calls by trimming low quality portions of the reads. Our simulation results on both randomly generated reads where methylation status is known and real world data indicate that ME-plot can detect errors in methylation mapping procedures and suggest post-processing steps to reduce the errors. ME-Plot requires SAM/BAM files for its analysis. **Availability:** The python package is available at the following URL. https://github.com/joshuabhk/methylsuite

## 1 INTRODUCTION

Whole genome bisulfite sequencing (WGBS) has been contributed to provide single nucleotide resolution DNA methylation throughout entire genome (Lister *et al*., 2009). WGBS deepens our understanding of DNA methylation in key biological processes such as genomic imprinting, transcriptional regulation and silencing of transposable elements. As a high throughput measurement, WGBS poses a novel problem of mapping bisulfite-treated reads, since the bisulfite treatment changes unmethylated cytosines into thymines; to address this, several mapping approaches has been developed (Krueger and Andrews, 2011; Xi and Li, 2009; Hansen *et al*., 2012).

As in other next-generation sequencing techniques, quality control of the bisulfite treated sequencing reads is important for accurate measurements. The M-bias plot has been developed as the first bisulfite sequencing quality control tool (Hansen *et al*., 2012) and it is also used in BSeQC package (Lin *et al*., 2013). M-bias plots show CpG methylation rates in each position in the reads when all reads are aligned to their 5’ ends. If there are systematic biases during sequencing procedures such as 3’ overhang end repair, 5’ bisulfite conversion failure and 3’ low quality sequences (Bock, 2012), the methylation rates should differ between read positions. Based on this assumption, M-bias plot is able to identify biased read positions. Another bisulfite sequencing quality control tool is Bis-SNP (Liu *et al*., 2012). As part of SNP caller using bisulfite sequencing data, Bis-SNP corrects 5’ bisulfite conversion failures. However, M-bias plot cannot differentiate the systematic bias introduced due to the bisulfite sequencing versus errors introduced due to mapping errors or low quality sequences.

## 2 METHODS

ME-plot uses pileup of mapped reads in SAM/BAM like M-bias plot. However unlike the M-bias plot, ME-plot additionally reports bisulfite specific errors in each methylation position in the reads. For each context of methylation (CpG, CHG and CHH, where H is a nucleotide other than G), ME-plot reports its average rates of mismatches, i.e. rate of A or G matched to C. In addition, ME-plot reports average methylation rates for all methylation contexts like the M-bias plot.

ME-plot also contains a cytosine methylation rate caller program that takes input from SAM/BAM mapping files. All trimming parameters determined from ME-plot are readily applicable to the methylation caller. Additionally, since our methylation caller can accept de facto standard mapping formats like SAM/BAM input files to produce the methylation rates, it can be readily used other bisulfite sequencing mappers without calling scripts like GSNAP (Wu and Nacu, 2010) for single end reads generated by MethylC-Seq protocol (Lister *et al*., 2009). We also introduced a formula to obtain the corrected amount of cytosine in our methylation caller, since the thymine in the bisulfite sequencing can be from bisulfite converted cytosine or from genomic thymine.

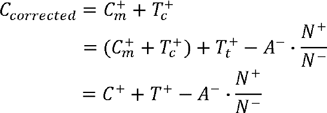

Where, 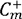 is the number of methylated cytosines which is same as observed cytosines or 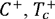 is the number of thymines converted from unmethylated cytosines (cannot be directly measured), 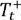is the number of thymines from thymine SNP (*T*^+^is the actual number of observed thymines, so 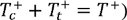is the number of adenocines from the other strand, *N^+^*is the sequencing depth on the strand with the cytosine, *N*^−^ is the sequencing depth on the other strand. The formula assumes 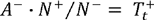since the ratio between *N^+^* and *N*^−^ compensates the possible sequencing depth difference between the two strands.

Random reads were generated from hg19 chromosome 21 sequences. After SNP introduction (based on dbSNP common SNPs) and sequencing errors were simulated according to the Phred score associated with GSM675542. Random reads were generated in various sizes 1M, 5M, 10M and 20M to track the changes of errors and coverage according to the input sizes. We developed and tested our code in python 2.7 (it is compatible with python 2.5 or above). All simulation code and data can be found at the same URL as ME-plot package.

## 3 RESULTS

We demonstrated the usage of ME-plot on two datasets: 1) simulated random reads dataset from hg19 chromosome 21, 2) WGBS from H1ESC (GSM675542). All reads were mapped both using Bismark and BSMAP and methylation was called by ME-plot methyl-caller. Our simulation results indicate that the average error per cytosine is higher if reads are not trimmed by quality scores when the sequencing quality scores accurately reflect the errors from sequencing (Fig. 1a). Conversely, if the reads are trimmed by quality score 20 after mapping, the reads are trimmed too much and the average error is higher than trimmed by quality score 10 or 4. In our simulation the quality score of 4 achieves a balance between of read coverage and pruning bad quality 3’ ends of sequencing reads. Interestingly our correction of measuring T on the opposite strand (Trim4/AllNT category in Fig. 1a) shows further accuracy improvement presumably by correcting the remaining mismatches in the mapping results. The results are consistent for datasets with different amounts of reads (1M, 5M and 10M) and for both bisulfite read mappers. The additional results are presented in the supplementary material.

**Figure 1.**
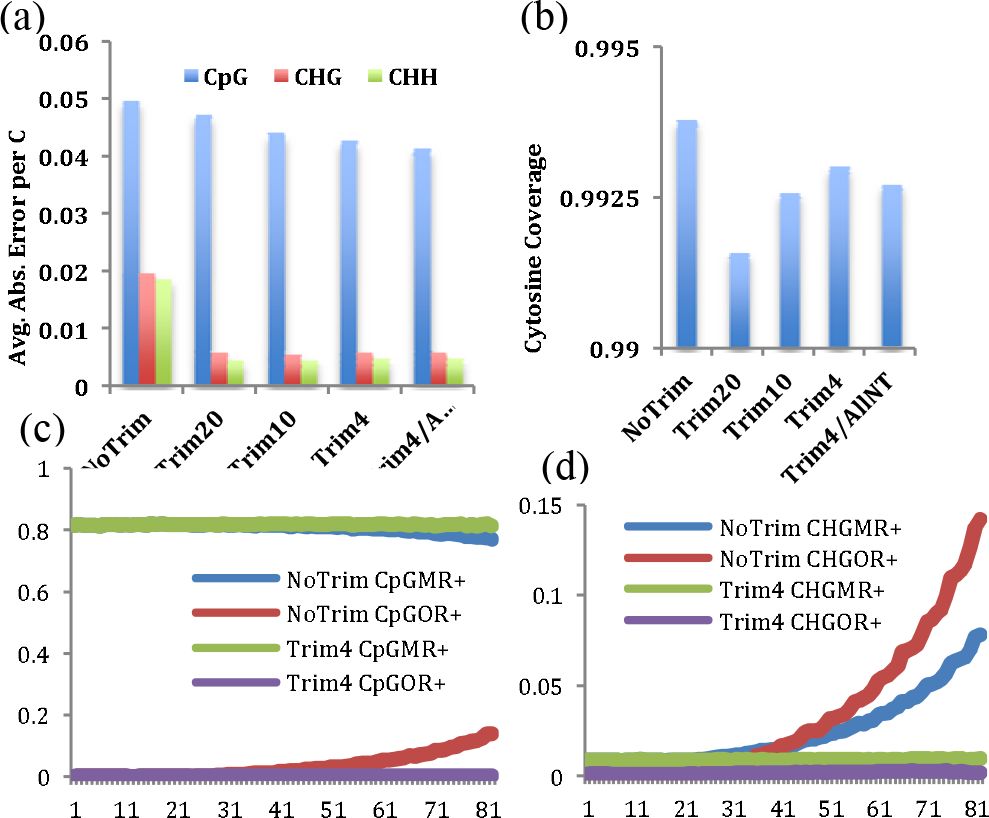
Relationship between methylation calling and M-bias plot and ME-Plot on a simulation dataset. (a) Bismark methylation calling performances on reads that were not trimmed (NoTrim), post-mapping trimming by Phred quality score 20, 10 and 4 (Trim20, Trim10 and Trim4) respectively. Trim4/AllNT is methylation rates after further correction (see methods). (b) The changes of cytosine coverage according to the trimming methods. (c) The ME-plot results of average methylation rates of each position in the mapped reads on CpGs for no trimmed dataset (NoTrim CpGMR+), trimmed by Phread quality score 4 (Trim4 CpGMR+) and error rates on CpGs for no trimmed dataset (NoTrim CpGOR+), and for trimmed by Phred quality score 4 (Trim4CpGOR+). (d) Similar to (c) but on CHG.

The ME-plot indicates the cause of the lower accuracies in the no-trimmed methylation calling results (Fig. 1c and d). Since ME-plot summarizes the mismatch rates at each methylation sites in CpG, CHG, and CHH contexts and reports the average mismatch rates in each position in the reads, mismatch rates of the no-trimmed dataset (NoTrim CpGOR+ and NoTrim CHGOR+ in Fig 1.c and d) highlights that the mapping errors are increasing toward the 3’ end of the reads. The CpG methylation rate of no-trimmed dataset in Fig 1c shows a slight decrease in the methylation rates due to the random C to T sequencing error and T to C error would increase the methylation rates in CHG or CHH contexts as shown in Fig 1d since they are generally not methylated.

Since there is no gold standard for methylation rates in the real dataset from H1ESC, we used consistency between the two repli-cates as a measure of our quality trimming, as in a previous study (Lin *et al*., 2013). As expected, the agreement between the methylation rates between the two replicates, became higher after the quality control. We used Kullback-Leibler distance to measure the differences between the two distributions and the distances were 0.0705 before QC and it dropped half to 0.032 after the postprocessing. Interestingly, the real dataset revealed an additional benefit of ME-plot, indicating bias in the last two positions that is difficult detect in the M-bias plot (Supplemental figure).

## 4 CONCLUDING REMARKS

It is worth noting that methylation rates in M-bias plots do not have much meaning in the absolute sense, rather the importance is in the relative methylation rates against the read positions. In other words, a better dataset results in a more horizontal straight line regardless of the methylation values. However, ME-plot’s error rates have a meaning both in the absolute and relative sense unlike the M-bias plot. Like the M-bias plot, the changing error rate according to the read position indicates biases in the read positions. Also, generally higher error rates (without any changes according to the read position) might indicate some systemic low quality of the datasets or some problems in the mapping procedures or parameters. We demonstrated that ME-plot can serve as a comple-ment QC tool to the existing methods by using simulated datasets and a real dataset. A generalized methylation caller accompanied by ME-plot can be also a useful tool for methylation calling proce-dures and provides additional benefits by correcting the mapping errors or SNPs. In summary, we believe ME-plot can facilitate analysis of DNA methylation and help development of other tools in methylation analyses.

## ACKNOWLEDGEMENTS

The authors are grateful to Brian Pierce for his helpful comments and critical reading of the manuscript.

